# Polymorphic mobile element insertions contribute to gene expression and alternative splicing in human tissues

**DOI:** 10.1101/2020.05.23.111310

**Authors:** Xiaolong Cao, Yeting Zhang, Lindsay M Payer, Hannah Lords, Jared P Steranka, Kathleen H Burns, Jinchuan Xing

## Abstract

**Background:** Mobile elements are a major source of human structural variants and some mobile elements can regulate gene expression and alternative splicing. However, the impact of polymorphic mobile element insertions (pMEIs) on gene expression and splicing in diverse human tissues has not been thoroughly studied. The multi-tissue gene expression and whole genome sequencing data generated by the Genotype-Tissue Expression (GTEx) project provide a great opportunity to systematic determine pMEIs’ role in gene expression regulation in human tissues.

**Results:** Using the GTEx whole genome sequencing data, we identified 20,545 high-quality pMEIs from 639 individuals. We then identified pMEI-associated expression quantitative trait loci (eQTLs) and splicing quantitative trait loci (sQTLs) in 48 tissues by joint analysis of variants including pMEIs, single-nucleotide polymorphisms, and insertions/deletions. pMEIs were predicted to be the potential causal variant for 3,522 of the 30,147 significant eQTLs, and 3,717 of the 21,529 significant sQTLs. The pMEIs associated eQTLs and sQTLs show high level of tissue-specificity, and the pMEIs were enriched in the proximity of affected genes and in regulatory elements. Using reporter assays, we confirmed that several pMEIs associated with eQTLs and sQTLs can alter gene expression levels and isoform proportions.

**Conclusion:** Overall, our study shows that pMEIs are associated with thousands of gene expression and splicing variations in different tissues, and pMEIs could have a significant role in regulating tissue-specific gene expression/splicing. Detailed mechanisms for pMEI’s role in gene regulation in different tissues will be an important direction for future human genomic studies.

## Introduction

Mobile genetic elements, or mobile elements (MEs), are segments of DNA that can move around and make copies of themselves within a genome [1]. At least 50% of the human genome is derived from MEs [2] and three non-long terminal repeat (non-LTR) retrotransposons dominate the recent ME activity: the short interspersed element (SINE) *Alu* [3], the long interspersed element 1 (LINE1) [4], and the composite SVA (SINE-VNTR (variable-number tandem repeat)-*Alu*) [5, 6]. LINE1 is an autonomous ME and encodes proteins that are required for the retrotransposition of itself [7], non-autonomous *Alu* and SVA elements [8], as well as occasionally cellular RNAs [9]. Many diseases, including cancer [10] and psychiatric disorders [11], are associated with the activities of MEs [12, 13]. In addition to causing genomic structural changes, MEs can also alter mRNA splicing [14] and gene expression levels [15, 16] via a wide variety of mechanisms, including acting as promoters [17], enhancers [18], splicing-sites [19], terminators for transcription [20], and affecting chromatin looping [21].

The activities of MEs create insertional mutations and other structural rearrangement of genomic DNA, leading to thousands of polymorphisms among human individuals and populations [22-24]. The effect of polymorphic mobile element insertions (pMEIs) on gene expression have been studied in the 1000 Genomes Project (1KGP) samples [25-28] and human induced pluripotent stem cells [28]. Using the gene expression data from the transformed B-lymphocytes cell lines of the 1KGP samples and iPSCs, several hundred pMEI loci were identified as expression quantitative trait loci (eQTLs). However, the extent of pMEIs’ impact on human gene expression in diverse tissues has not been extensively examined.

The Genotype-Tissue Expression (GTEx) project aims at building a public resource to study tissue-specific gene expression and regulation [29-31]. The genetic associations for gene expression and splicing were analyzed as the main topic of the project. In the v7 release, GTEx provides 11,668 high-depth RNA-sequencing (RNA-seq) data from 51 tissues and 2 cell lines of 714 donors. More than 600 of the donors have also been subjected to high-depth whole genome sequencing (WGS). This rich dataset makes it possible to assess the impact of different types of genomic variants on gene expression, such as structural variants [32], rare variants [33], and short tandem repeats [34]. However, the role of pMEIs in gene regulation and alternative splicing, especially pMEIs that are not present in the reference genome, have not been fully evaluated. Given thousands of common pMEIs are present in human populations, they might represent a major type of variants associated with gene expression regulations. With the large GTEx data set, we systematically identified pMEIs in each donor, and examined the impact of common pMEIs on gene expression and splicing.

## Result

### Detection of pMEIs in GTEx individuals

We obtained WGS data from the GTEx v7 release. Using MELT [35], we identified MEs that are present in the sequenced individuals but absent in the reference genome, as well as MEs that are present in the reference genome but absent in a subset of sequenced individuals. We refer to these two types of ME polymorphisms as nonreference MEIs (nrMEIs) and reference MEIs (rMEIs) in the following text, respectively. After filtering, we identified a total of 80,057 candidate nrMEIs and rMEIs in 639 individuals, including 638 GTEx individuals and the HuRef sample (Table 1). Overall, 99.5% of sites have no-call rates < 25%, demonstrating the high quality of the sequenced genomes.

**Table 1:** Overview of pMEIs in the MELT callset, eQTL, and sQTL analyses.

| ME type | MELT callset |  |  | eQTL |  |  | sQTL |  |  |
| --- | --- | --- | --- | --- | --- | --- | --- | --- | --- |
|  | raw | HQ | common | all | causal | highest | all | causal | highest |
| nrAlu | 62,864 | 13,870 | 2,157 | 1,451 | 562 | 147 | 1,071 | 539 | 191 |
| nrL1 | 11,159 | 2,130 | 246 | 177 | 81 | 23 | 126 | 71 | 18 |
| nrSVA | 1,877 | 558 | 69 | 61 | 32 | 12 | 51 | 27 | 13 |
| rAlu | 3,837 | 3,687 | 968 | 671 | 253 | 84 | 444 | 202 | 106 |
| rL1 | 192 | 188 | 59 | 42 | 15 | 7 | 28 | 18 | 8 |
| rSVA | 128 | 112 | 21 | 20 | 13 | 9 | 14 | 9 | 6 |
| **Total** | **80,057** | **20,545** | **3,520** | **2,422** | **956** | **282** | **1,734** | **866** | **342** |

MELT callset: raw: all pMEI loci identified by MELT; HQ: high quality loci after quality control; common: pMEIs used for eQTL and sQTL analysis.
eQTL/sQTL analysis: all: unique pMEIs in eQTL/sQTL analysis (FDR<10%); causal: unique pMEIs identified as the causal variant; highest: unique pMEIs identified as the causal variant with highest causal probability.

The raw ME loci were filtered based on quality scores, no-call rates, and other criteria (see methods for detail). After filtering, 20, 545 high quality loci were selected for further analysis. Most pMEIs have allele frequency less than 0.05, especially nrMEIs (Fig. S1a, S1b). Because the human reference genome is based on only a small number of individuals, pMEIs present in the reference genome (rMEIs) should be more common than pMEIs absent in the reference genome (nrMEIs). As expected, overall rMEIs have higher allele frequencies than nrMEIs (Fig. S1a, S1b). The numbers of pMEI loci in different individuals are correlated with the self-reported ancestry of the individuals. In general, the number of nrMEI and rMEI loci in African individuals is larger than in non-Afircan individuals (Fig. S1c). We define common pMEIs as those with allele frequency between 0.05 and 0.95. Overall, 3,076 nrMEIs and 1,662 rMEIs are common, which are 18.58% and 41.68% of the high quality nrMEI and rMEI call sets, respectively. After further quality control, a total of 3,520 common pMEIs were selected for the following analyses (Table 1).

### Identify pMEI-associated eQTLs

Next, we determined the effect of pMEIs on nearby gene expression by mapping pMEI-associated cis-eQTLs. The GTEx v7 release includes expression data of 56,202 genes from 53 tissues/cell lines, with 19,820 protein-coding genes and 36,382 non-coding genes (Table 2). We selected 46 tissues and 2 cell lines with expression data in more than 70 individuals for the analysis (ranging from 78 to 481 individuals per tissue/cell line) (Table S1). We will refer both tissues and cell lines as tissues for simplicity in the following text. After excluding low-expressed genes from the analysis, the average number of tested coding genes in each tissue is 16,461 with a standard deviation (SD) of 598 (see methods for detail). For non-coding genes, the testis is an outlier with 14,970 expressed genes. The average number of expressed non-coding genes in tissues other than testis is 7,294 with an SD of 826.

**Table 2:** Summary of genes.

| Gene | Total | Expressed | eQTLs | ME causal | ME highest causal |
| --- | --- | --- | --- | --- | --- |
| protein-coding | 19,820 | 19,064 | 4,243 | 1,062 | 294 |
| non-coding | 36,382 | 19,111 | 2,099 | 526 | 139 |
| **Total** | **56,202** | **38,175** | **6,342** | **1,588** | **433** |

Expressed: genes used in eQTL analysis of at least one tissue.
eQTLs: number of unique genes in ME-only eQTL analysis with FDR < 10%
ME causal, ME highest causal: unique genes with pMEIs predicted as causal variant or causal variant with highest probability, respectively

We performed cis-eQTL mapping with Matrix eQTL [36] in each tissue. Here we define an eQTL as a unique combination of tissue-gene-variant. Among all tissues, we identified 30,147 eQTLs with 6,342 distinct genes, 2,422 distinct pMEIs, and 8,204 distinct gene-ME pairs with a false discovery rate (FDR)<10%. pMEIs that are eQTLs showed strong enrichment near the transcription start site (TSS) of genes, although some eQTLs-pMEIs are much further away from the associated genes (Fig. S2). Next, we define an eGene as a tissue-gene pair that were identified in the eQTL analysis with an FDR<10%, while an eVariant as a tissue-variant pair with an FDR<10%. Because an eGene can be influenced by multiple variants and an eVariant may have impact on multiple genes, the numbers of eGenes (24,109) and eVariants (17,230) are smaller than the total number of eQTLs. The number of eQTLs (FDR < 10%) per tissue ranges from 118 to 1,609 and the sample size is strongly correlated with the number of detected eQTLs (r^2^ = 0.85, Fig. 1a, 1c, Table S1). This strong correlation between the number of detected eQTLs and the sample size was also observed in similar studies [30, 31, 34]. The correlation is even stronger (r^2^ = 0.92) when we added the number of expressed genes as a covariate in the linear regression analysis of the number of eQTLs. For eQTLs, most gene-ME pairs were identified in only one tissue, account for 53% of coding genes and 62% of noncoding genes (Fig. 1d, Table S1). The higher tissue-specificity of noncoding gene eQTLs could be explained by the fact that noncoding genes more frequently have tissue-specific expression patterns.

**Fig. 1.**
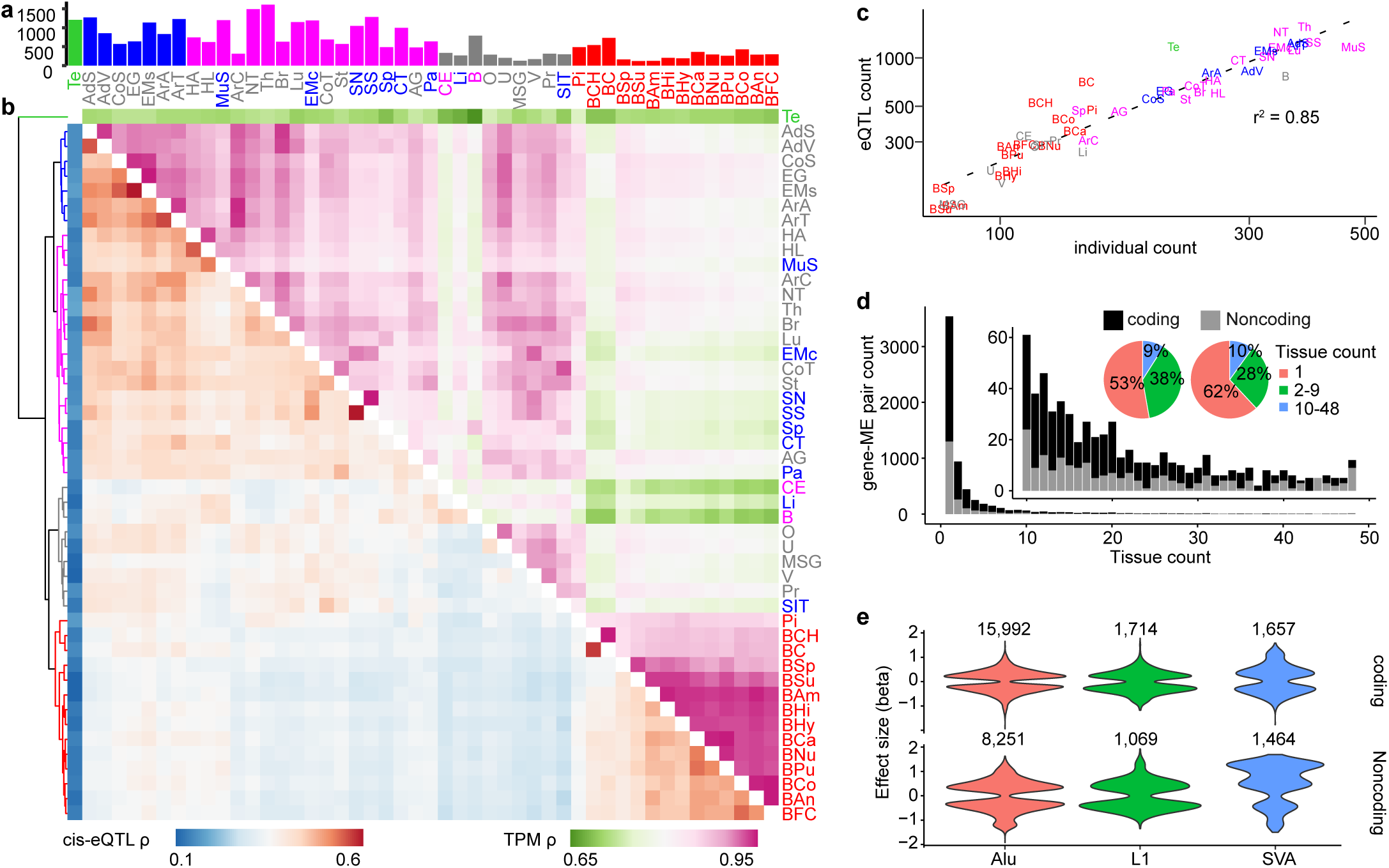
Overview of the ME-only eQTL analysis. (**a**) The number of detected eQTLs with Benjamini-Hochberg FDR < 10% in each tissue. Bars are colored by tissue clusters based on cis-eQTL as shown in (**b**, tree). (**b**) Similarity (Spearman’s correlation coefficient ρ) between different tissues based on cis-eQTL FDR values (lower triangle) and gene expression TPM values (upper triangle). Gene-pMEI pairs with FDR < 10% in at least one tissue is selected for the analysis. Tree on the left of the plot was based on the hierarchical clustering of the cis-eQTL results and the branches are colored to five groups. Tissue text colors in (**a, b**) were based on hierarchical clustering tree of TPM results (data not shown). (**c**) The relationship between the eQTL count (FDR < 10%) and the individual count in different tissues. Tissue text is colored by tissue clusters based on cis-eQTL in (**b**, tree). The axes are in log scale. (**d**) Gene-pMEI pair count and the number of tissues they were detected as significant for coding and noncoding genes. (**e**) Effect size (beta value) distribution for coding and noncoding eQTLs of different types of pMEIs. Tissue abbreviations: AdS, Adipose Subcutaneous; AdV, Adipose Visceral Omentum; AG, Adrenal Gland; ArA, Artery Aorta; ArC, Artery Coronary; ArT, Artery Tibial; BAm, Brain Amygdala; BAn, Brain Anterior cingulate cortex BA24; BCa, Brain Caudate basal ganglia; BCH, Brain Cerebellar Hemisphere; BC, Brain Cerebellum; BCo, Brain Cortex; BFC, Brain Frontal Cortex BA9; BHi, Brain Hippocampus; BHy, Brain Hypothalamus; BNu, Brain Nucleus accumbens basal ganglia; BPu, Brain Putamen basal ganglia; BSp, Brain Spinal cord cervical c-1; BSu, Brain Substantia nigra; Br, Breast Mammary Tissue; CE, Cells EBV-transformed lymphocytes; CT, Cells Transformed fibroblasts; CoS, Colon Sigmoid; CoT, Colon Transverse; EG, Esophagus Gastroesophageal Junction; EMc, Esophagus Mucosa; EMs, Esophagus Muscularis; HA, Heart Atrial Appendage; HL, Heart Left Ventricle; Li, Liver; Lu, Lung; MSG, Minor Salivary Gland; MuS, Muscle Skeletal; NT, Nerve Tibial; O, Ovary; Pa, Pancreas; Pi, Pituitary; Pr, Prostate; SN, Skin Not Sun Exposed Suprapubic; SS, Skin Sun Exposed Lower leg; SIT, Small Intestine Terminal Ileum; Sp, Spleen; St, Stomach; Te, Testis; Th, Thyroid; U, Uterus; V, Vagina; B, Whole Blood.

To determine if closely related tissues show similar eQTL profiles, we evaluated the eQTL correlations among different tissues for ME-gene pairs using Spearman’s correlation (ρ) (Fig. 1b). The Spearman’s correlation of the expression level of these eQTL genes (calculated as Transcript per million (TPM)) were also calculated to determine the impact of similarities of gene expression on eQTL identification. As shown in Fig. 1b, tissues from different brain regions were clustered together by eQTL correlations and eQTL gene expression levels. Testis (Te) showed the highest difference with other tissues in both eQTLs and gene expression levels. Highly similar tissues, such as skin sun exposed (SS) and skin not sun exposed (SN), brain cerebellum (BC) and brain cerebellum hemisphere (BCH), are highly similar in the eQTL significance (ρ > 0.6) and the gene expression level (ρ > 0.95). However, whole blood and EBV-transformed lymphocytes (B and CE) showed lower gene expression correlation with other tissues (ρ < 0.8 in general) than other tissue pairs, suggesting a different expression pattern in blood and cell line samples. It is also obvious that the correlations are higher for gene expression than for eQTLs. This is partially because gene expression values can be more accurately determined and normalized than eQTL significance values.

To determine if the presence/absence of a pMEI has a directional impact on the gene expression, we examined the effect size and direction (positive or negative) of different types of pMEIs. We observed no statistically significant difference in the direction and the scale of the effect sizes between the presence versus the absence of pMEIs, for all three types of pMEIs, and for both coding and non-coding genes (Fig. 1e). This result suggests that for common pMEIs, ME-specific sequence feature is a less important factor affecting the nearby gene expression than the presence/absence of a pMEI. We also compared the correlation of the direction of the effect (i.e., the sign of the beta value) among tissue pairs. Overall, the direction of effect for pMEIs are highly consistent among tissue pairs, with an apparent exception of testis (Fig. S3). Excluding testis, the effect direction among tissue pairs are consistent for 98.6 ± 1.7% of eQTLs.

### Fine-mapping causal pMEIs for eGenes

Due to the linkage disequilibrium among genetic variants, a number of tightly linked variants can be identified as eQTLs along with the causal variant. To determine whether the pMEIs identified in the eQTL analysis are the causal variants, we applied a fine mapping approach for each eQTL locus. To do this, we gathered the single nucleotide polymorphisms (SNPs) and insertions/deletions (indels) from GTEx individuals and selected a total of 6,334,405 high quality common variants, including 5,837,891 SNPs and 496,514 indels. For the 6,342 unique genes identified in the ME-only eQTL analysis, we performed joint analyses for pMEIs and these common variants to identify all variants associated with an eGene in each tissue. Then, we applied fine-mapping method for each of the 24,109 eGenes to identify the contributions of MEs in altering gene expression. Overall, pMEIs were included in the causal variant set for 13.98% of eGenes, ranging from 10.69% in Sun Exposed Lower Leg Skin to 25.33% in Hippocampus among tissues (Table 1, Table S1). pMEIs were detected as the highest-probability causal variant for 4.55% of tested eGenes (2.67 – 9.18% among tissues), slightly larger than 3.5% (2.4% – 4.4% among tissues) of general structural variants in a previous study [32] (Table 1, Table S1).

### Enrichment of eQTL-pMEIs in functional genomic elements

To explore the potential molecular mechanisms for pMEIs’ influence on gene expression, we determined the enrichment of pMEIs in functional genomic elements. We grouped the 3,520 common pMEIs to three categories: not an eVariant for any gene (NS), identified as an eVariant but not a causal eVariant (Related), and identified as a causal eVariant (Causal). Compare to the NS set, pMEIs that are eVariants (Related and Causal) are significantly enriched in enhancers, 10 kb upstream or downstream the gene, and exons and introns of affected gene (Fig. 2a-e). This observation is consistent with the observation that eQTL-pMEIs are enriched near the TSS of genes (Fig. S2). Importantly, pMEIs in the “Causal” category are more enriched in functional regions than “Related” pMEIs in all categories except in introns. This enrichment suggests pMEIs in the casual set are more likely to be the true functional variant for the gene expression change. Only a small portion of pMEIs are in the exon of genes, and all of them are detected as eQTLs and showed a stronger enrichment in the causal set (Fig. 2e). Given the size of the pMEIs, it is expected that the exonic pMEIs will have a strong impact on the gene expression level. The enrichment of pMEIs in functional elements are similar to structural variants in general, as structural variants impacting gene expression are also enriched in enhancers, promoters, and regions close to the affected genes [32].

**Fig 2.**
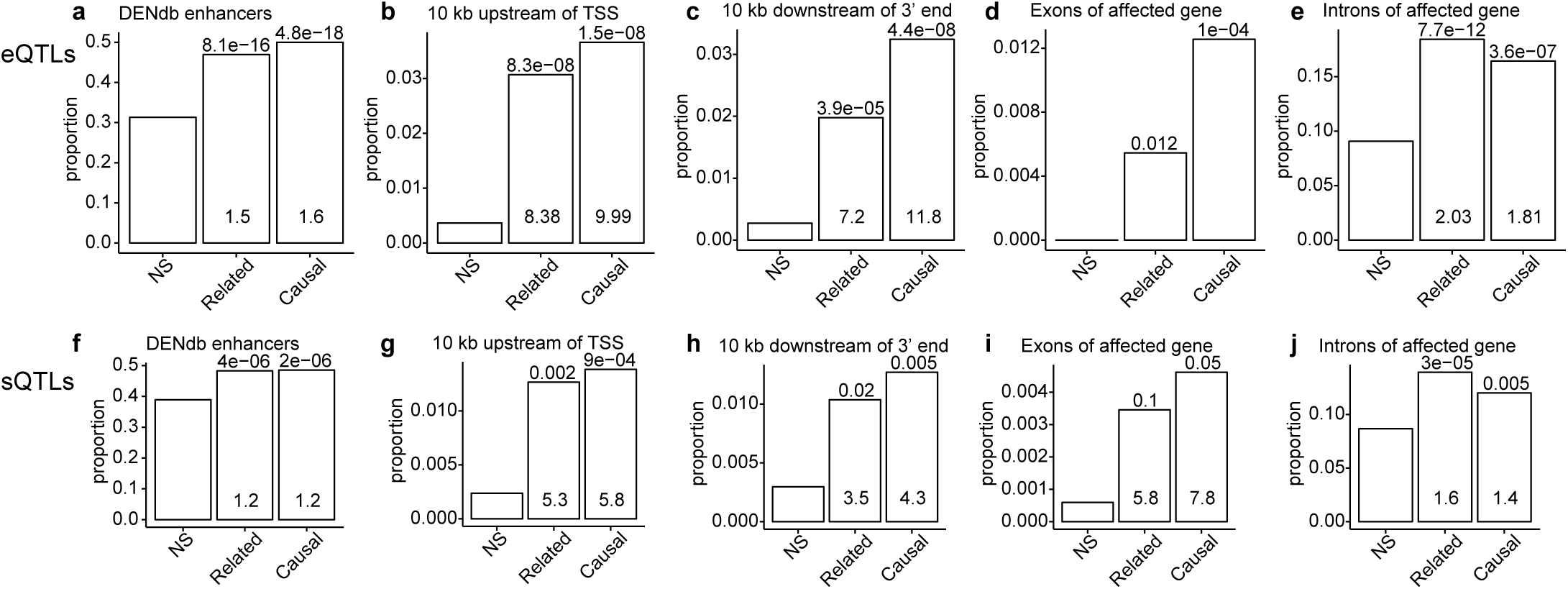
Enrichment of pMEIs in different functional genomic regions of affected genes in eQTL analysis (**a** - **e**) and sQTL analysis (**f** - **j**). Functional genomic regions include enhancers from the Dragon Enhancers Database (DENdb) (**a, f**); 10 kb upstream from the transcription starting site (TSS) (**b, h**), 10 kb downstream (**c, i**), exons (**d, j**), and introns of the affected gene (**e, k**). pMEIs were divided into three categories: NS, pMEIs that were not reported to be significantly related with any gene or ASE in any tissue; Related, pMEIs that were significantly associated with at least one gene or ASE, but were not reported as causal; Causal, pMEIs that were reported as causal for at least one gene or ASE (see methods for detail). Bar plot shows the proportion of pMEIs in each genomic feature in each category (NS, Related, or Causal). Values inside the bars are fold enrichment compared to NS, and values above the bars are p-value from Fisher’s exact test for significance of enrichment compared to NS. For exons in eQTL analysis in (**d**), the fold enrichment values are not available as the proportion of pMEIs in exon is zero in NS.

### Identify pMEIs-associated sQTLs

We next investigated the impact of pMEIs on alternative splicing of genes. We performed the analysis of splicing quantitively trait loci (sQTLs) similar to eQTLs, using PSI (percent splicing in) scores of alternative splicing events (ASEs) instead of TPM of genes (see method for a full definition of the ASEs). When determining ASEs, genes sharing one or more exons were grouped together as a gene cluster. We will refer to these gene clusters as genes in the sQTL analysis for simplicity. There are 165,882 ASEs from 17,015 genes (Table 3). About half of the events occur inside the gene, including alternative 3’/5’ splicing site (A3/A5), mutually exclusive exons (MX), retained intron (RI), and skipped exon (SE). The other half occur at the edge of a gene, including alternative first/last exons (AF/AL) (Table 3). We detected a total of 21,529 sQTLs with 7,184 distinct splicing events from 2,992 genes with FDR < 10%, among which 11,183 and 10,346 sQTLs are from events internal and at the edge of the gene, respectively. The numbers of detected sQTLs with events internal or at the gene edge are proportional to the total number of possible events in these regions, indicating weak or no selective preference. The number of sQTLs in each tissue ranges from 81 to 1,120 (Fig. 3a, Table S2). Similar to eQTLs, the number of sQTLs is highly correlated with the number of donors for each tissue (r^2^ = 0.71, Fig. 3c). Also, 83% internal and 73% of gene edge splicing event-pMEI pairs are only detected in one tissue, suggesting the impact of pMEIs on gene splicing is highly tissue-specific (Fig. 3d). However, sQTL analysis uses the transcript level PSI information, which is noisier than the gene level TPM used in the eQTL analysis. Therefore, the higher tissue-specificity of sQTLs than eQTLs may also partly due to the lower power and higher level of false-negatives in the sQTL analysis. Although sQTLs appear highly tissue-specific, we did identify similarities among related tissues (e.g., brain regions) based on sQTL significance and PSI scores for ASEs (Fig. 3b), and high agreement in the direction of pMEIs’ effect (Fig. S3), similar to eQTLs. Overall, tissues show more variance based on gene alternative splicing (PSI values) than gene expression levels (TPM values), and the similarity of sQTL and PSI metrics are less than eQTL and TPM metrics. The effect size for sQTLs can be either positive or negative (Fig. 3e), but values of beta are much smaller than eQTLs due to the small variation of PSI values (0-1).

**Table 3:**
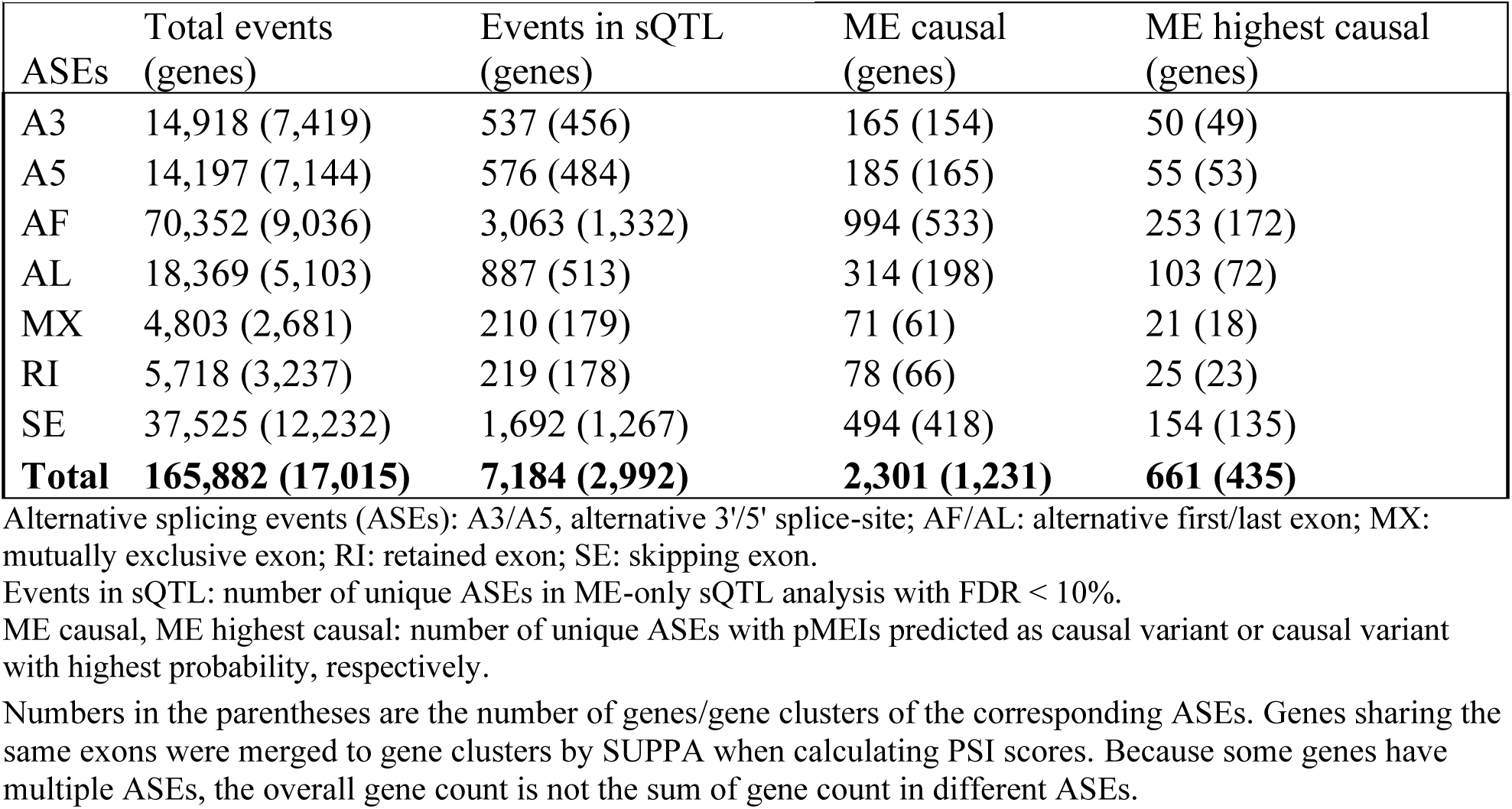
Summary of alternative splicing events.

| ASEs | Total events (genes) | Events in sQTL (genes) | ME causal (genes) | ME highest causal (genes) |
| --- | --- | --- | --- | --- |
| A3 | 14,918 (7,419) | 537 (456) | 165 (154) | 50 (49) |
| A5 | 14,197 (7,144) | 576 (484) | 185 (165) | 55 (53) |
| AF | 70,352 (9,036) | 3,063 (1,332) | 994 (533) | 253 (172) |
| AL | 18,369 (5,103) | 887 (513) | 314 (198) | 103 (72) |
| MX | 4,803 (2,681) | 210 (179) | 71 (61) | 21 (18) |
| RI | 5,718 (3,237) | 219 (178) | 78 (66) | 25 (23) |
| SE | 37,525 (12,232) | 1,692 (1,267) | 494 (418) | 154 (135) |
| **Total** | **165,882 (17,015)** | **7,184 (2,992)** | **2,301 (1,231)** | **661 (435)** |

Alternative splicing events (ASEs): A3/A5, alternative 3'/5' splice-site; AF/AL: alternative first/last exon; MX: mutually exclusive exon; RI: retained exon; SE: skipping exon.
Events in sQTL: number of unique ASEs in ME-only sQTL analysis with FDR < 10%.
ME causal, ME highest causal: number of unique ASEs with pMEIs predicted as causal variant or causal variant with highest probability, respectively.
Numbers in the parentheses are the number of genes/gene clusters of the corresponding AEs. Genes sharing the same exons were merged to gene clusters by SUPPA when calculating PSI scores. Because some genes have multiple AEs, the overall gene count is not the sum of gene count in different AEs.

**Fig. 3.**
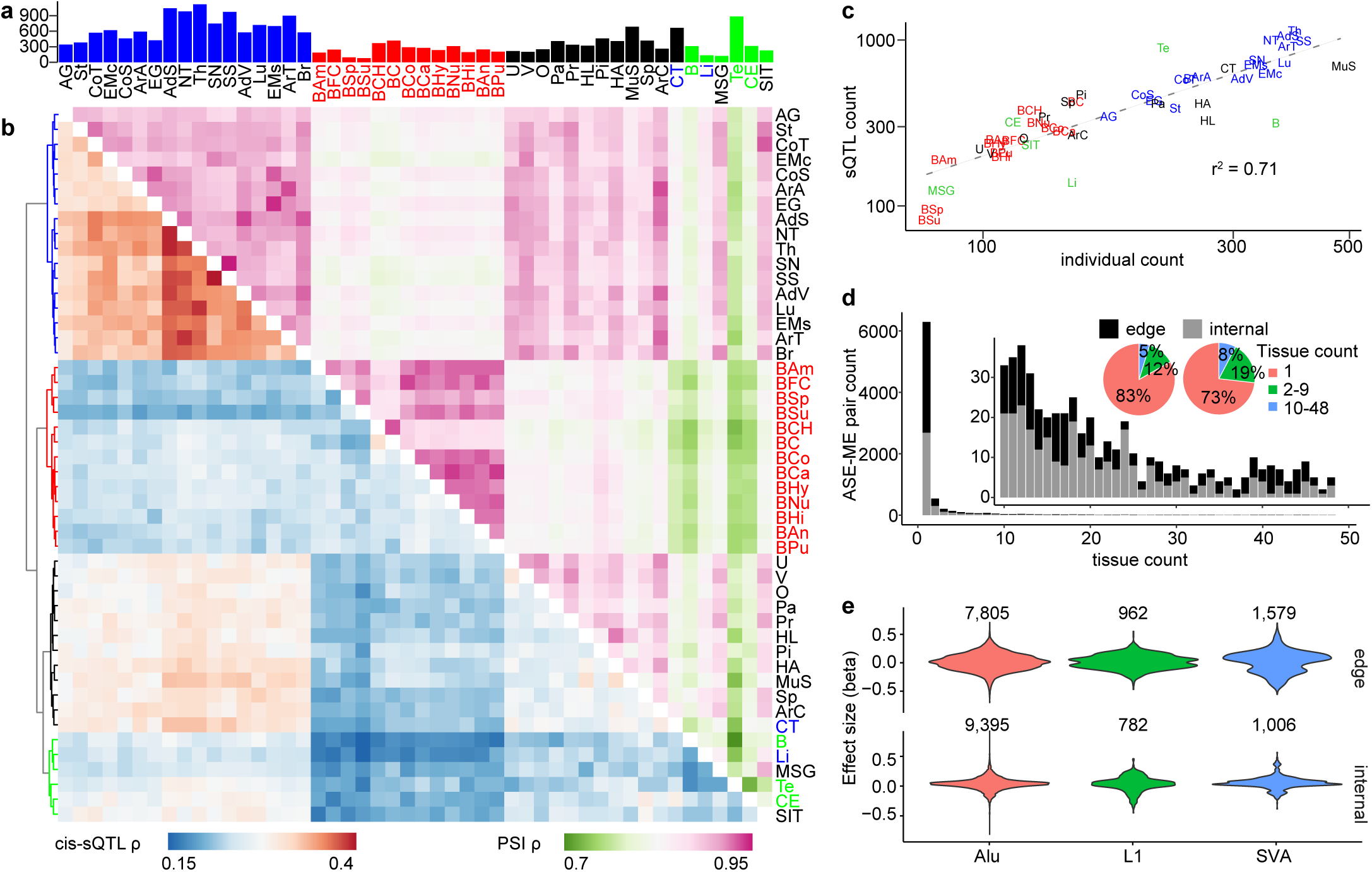
Overview of sQTL analysis in ME-only analysis. (**a**) Number of detected sQTLs with Benjamini-Hochberg FDR < 10% in each tissue. Bars are colored by tissue clusters based on cis-eQTL as shown in (**b**, tree). (**b**) Similarity (Spearman’s correlation coefficient ρ) between different tissues based on cis-sQTL (lower triangle) and alternative splicing events (ASEs) PSI values (upper triangle). ASE-pMEI pairs with FDR < 10% in at least one tissue is selected for the analysis. Tree was based on hierarchical clustering of the cis-sQTL results and the branches are colored to four groups. Tissue text colors in (**a, b**) were based on hierarchical clustering tree of PSI results (data not shown). (**c**) The relationship between the sQTL count (FDR < 10%) and the individual count in different tissues. The axes are in log scale. (**d**) ASE-pMEI pair count and the number of tissues they were detected as significant for events internal or at the edge of the gene. Tissue text is colored by tissue clusters based on cis-sQTL in (**b**, tree). (**e**) Effect size (beta values) distribution for ASEs internal or at the edge of different pMEIs. Tissue abbreviations are the same as in Fig. 1.

Next, we applied the fine mapping strategies to determine the causal pMEIs for sQTLs. pMEIs were identified as causal for 17.26% (13.10% – 35.11% among tissues) and as highest-probability causal for 4.33% (2.05% – 7.38%) of ASEs. Same as eQTLs, pMEIs detected as sQTLs (Related) or identified as causal variant for at least one ASEs (Causal) are significantly enriched in enhancer regions and regions close to the affected genes (Fig. 2f-i). However, the enrichment and significance of pMEIs are lower compared to eQTLs, likely because of the noisier measurement of PSI values than TPM values for eQTL analysis.

To determine if pMEIs affect the expression and splicing of genes simultaneously, we identified genes with both eQTLs and sQTLs. Both the significance and the effect size for eQTLs and sQTLs are positively correlated, indicating that a pMEI influences the expression of a gene is also likely to impact the alternative splicing and isoform abundance of that gene (Fig. 4a, b). Although ∼40% of pMEIs were identified in both eQTL and sQTL analysis, some pMEIs were only identified in one of the analysis, indicating different impact of pMEIs or different sensitivities of the two analyses (Fig. 4c). pMEIs detected only in sQTL analysis tend to have lower AF than pMEIs only in the eQTL analysis (Fig. 4d).

**Fig. 4.**
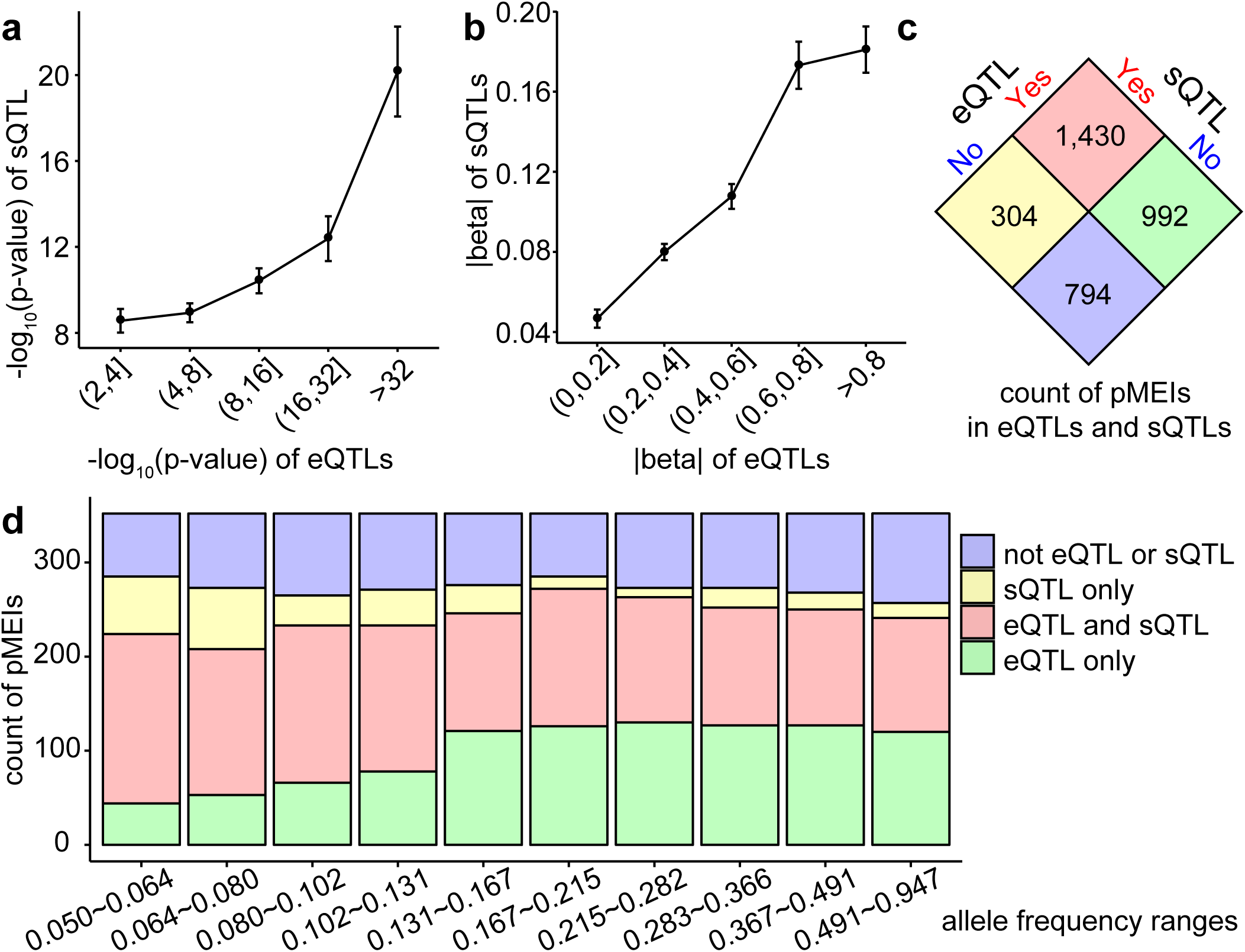
Correlation between eQTL analysis and sQTL analysis. (**a**) Correlation of p-values of eQTLs and sQTLs. Average −log10(p-values) of sQTLs were plotted against eQTLs with −log10(p-values) divided in five bins. (**b**) effect size (|beta|) of sQTL versus eQTL. Average |beta| of sQTLs were plotted against eQTLs with their |beta| values divided in five bins. (**a, b**) error bars are 95% confidence intervals. Only sQTL and eQTL pair that shared the same gene, tissue, and pMEI were included in the analysis. (**c**) Number of pMEIs detected in eQTL or sQTL analysis. **(d)** Count of pMEIs identified in eQTL or sQTL analysis in different allele frequency groups. The pMEIs were divided to 10 groups based on their allele frequencies so that each group have equal number of pMEIs.

### Experimental validation of eQTLs and sQTLs

To experimentally verify the predicted impact of specific pMEIs on gene expression and splicing, we evaluated selected loci in ectopic reporter assays (see methods for detail). We selected loci for validation based on the requirements of ectopic reporter assays (e.g., pMEI size, sequence availability, etc.), the supportive evidence from the eQTLs/sQTLs, and the importance of associated genes. For pMEIs predicted to be causal in the eQTL analysis, we selected six loci for experimental validation. All six tested ME loci showed significant difference in the gene expression between the presence and absence of the pMEI (p<0.01, unpaired 2-tailed t-test) (Fig. 5a). The presence of the pMEI resulted in upregulation of luciferase expression in five cases, with only one locus, *IP6K2*, where the presence of the pMEI reduced luciferase expression relative to the pre-insertion allele. These results indicate that pMEI in their genomic context can alter transcription levels, supporting their role as eQTLs. Three pMEIs have the same direction of effect (i.e. either up- or down-regulation in the presence of the pMEI) in the reporter assay as predicted computationally for the closest eGenes: *BDH2, PGR* and *IP6K2* (Fig. 5a). Because all three pMEIs are eQTLs in multiple tissues and in all tissues the pMEIs have the same predicted direction of effect, these pMEIs are likely to regulate gene expression across tissue types using a similar mechanism.

**Fig. 5.**
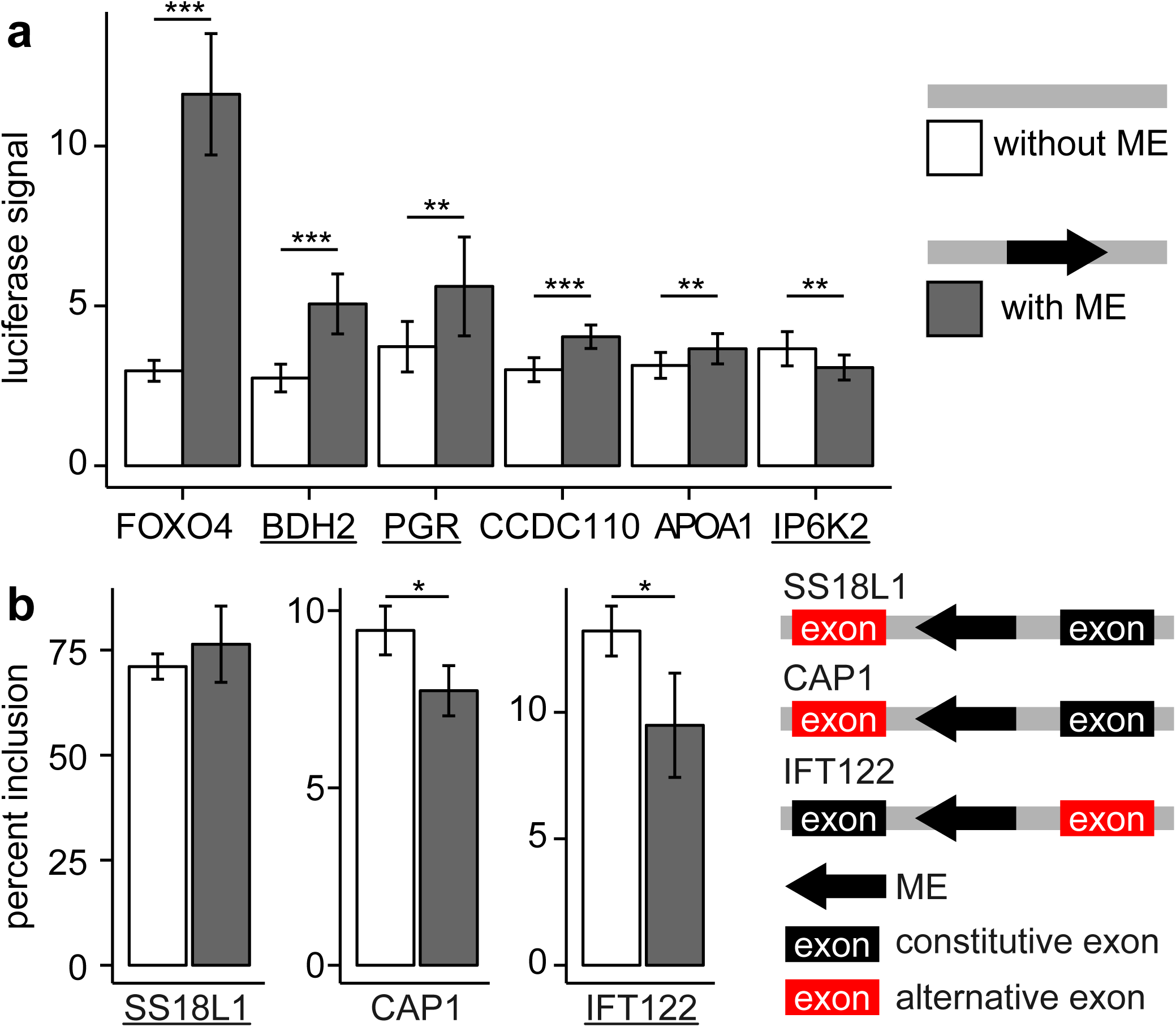
Experimental validation of eQTLs (**a**) and sQTLs (**b**). Gene names were labeled in x-axis, and those underlined showed effects in the same direction as predicted in computational analysis. For sQTL experiments, one constitutive exon was included with the alternative exon. Results are shown for the ME-containing construct and the construct without the ME. In (**b**), the direction of the arrow represents the strand of the ME on the chromosome. (* p < 0.05, ** p < 0.01, *** p < 0.001)

Next we performed experimental validation of pMEI sQTLs using ectopic reporter assays. We focused on pMEIs within genes and near differentially incorporated exons to enable evaluation with a minigene reporter. We evaluated three pMEI loci for sQTLs and identified significant effects of the ME at two of the three loci (p<0.05, unpaired 2-tailed t-test) (Fig. 5b). In both cases the presence of the pMEI resulted in less incorporation of the alternatively incorporated exon. We compared these results to the effects predicted for the pMEI-containing allele in our sQTL analysis. For *IFT122*, we predicted that the pMEI would decrease the exon inclusion in all tissues with sQTLs, and this prediction agrees with our ectopic assay. However, for *CAP1* the predicted effect of the presence of the ME on splicing did not agree with the experimental result. Altogether, these data confirm that pMEI can alter gene expression levels and isoform proportions largely consistent with the predicted effects in our QTL analysis.

## Discussion

MEs play important roles in gene regulation and have the capacity of creating new gene regulatory networks [18, 37, 38]. However, most previous studies on the impact of pMEIs on gene expression focused only on eQTLs and on lymphocyte cell lines (LCLs) from the 1KGP project [26, 32, 39]. The GTEx project provides an excellent opportunity to study the impact of pMEI on gene expression and alternative splicing in human tissues [30]. Although a previous study from the GTEx consortium included some pMEIs that are present in the reference genome (rMEIs), the study did not considered non-reference pMEIs and it is based on the smaller GTEx v6 release (147 individuals, 13 tissues) [32]. In this study we used MELT to identify pMEIs regardless of their annotation in the reference genome in more than 600 individuals. Combining the genotypes of common pMEIs with the GTEx RNA-Seq data, this dataset allows us to examine the impact of pMEIs on gene expression and gene splicing comprehensively in 48 tissues.

The high depth WGS from the GTEx project (mean coverage about 40-fold) resulted in sensitive pMEI identification and accurate genotyping [35]. We identified a total of 20,545 pMEI loci from 639 individuals, including 16,558 non-reference pMEIs and 3,987 reference pMEIs. The total number of pMEIs in our study is about ten times more than the 2,051 reference pMEIs identified in the previous study [32]. The number is also higher than the 17,934 pMEIs identified in the phase 3 of the 1KGP from 2,504 individuals, which was based on low-coverage WGS (mean coverage 7.4-fold) [27]. Only less than half of the pMEI loci (8,456) were identified both in this project and in the 1KGP. Recent studies showed that a large number of undiscovered pMEIs in individuals from different populations, especially for non-reference pMEIs [24, 40]. In addition, because of the repetitive nature of mobile elements, many pMEIs are missed by the current short-read sequencing technology [41]. Therefore, when more diverse populations are included and long-read sequencing technologies are used, we expect a lot more pMEIs will be identified.

We identified that the expression of 6,342 genes was correlated with 2,422 pMEIs and the splicing of 2,992 genes was correlated with 1,734 pMEIs in at least one of the 48 tested tissues. The number of pMEIs identified in eQTLs (2,422) in our study is much higher compared to previous studies [26, 28, 32, 39], such as 265 rMEIs reported in the GTEx SV study at FDR < 10% [32] and 235 pMEIs in an 1KGP study at FDR <= 5% [26]. We also identified a large number of pMEIs as the potential causal variant for eQTLs (956) and sQTLs (866). This difference highlights the value of both the large number of pMEIs identified from the high-coverage WGS data, and the many tissues examined in our study. The numbers of detected eQTLs and sQTLs in each tissue were highly correlated with the sample size of each tissue (r^2^ 0.85 and 0.71 for eQTLs and sQTLs, respectively) (Fig. 1c, Fig. 3c). Because the power of the QTL (eQTL and sQTL) analysis is closely related with the sample size, this linear relationship indicates that the sample size is still too small in most tissues. It is likely that many QTLs were not detected due to the small sample size in many tissues. In GTEx v8, there is no significant sign of eGenes/sGenes showing plateauing at sample size of 600 [31], suggesting more than 600 samples are needed to reach sufficient power to identify all eQTLs and sQTLs.

A previous study showed that difference analysis methods can produce very different eQTL results, even with the same raw dataset [28]. To assess the consistency of our eQTL analysis with other studies, we compared the eQTLs identified in LCLs with an eQTL study of LCLs from 1KGP samples [26]. Our data set contains 113 individual-derived LCLs, which is much smaller than the 445 LCLs in the 1KGP study [42]. With FDR < 5% as a cutoff, we identified 255 pMEI-associated eQTLs in GTEx LCLs. Despite that the differences in sequencing protocols, sample composition, and data processing, 67 of these eQTLs were also identified in the study of 1KGP eQTLs (Table S3). This result suggests that many of the pMEI-associated eQTLs are strong eQTLs that show consistent signal in individuals from different populations.

The significance of pMEI-associated eQTLs and sQTLs are similar in related tissues (Fig. 1b, Fig. 3b). Except for testis, tissue pairs also show strong consistency in the direction of pMEI’s effect in eQTLs and sQTLs (Fig. S3). Our results agree with a previous study showing that testis is unique in gene expression compared to other tissues [43]. The overall high consistency of eQTLs’ and sQTLs’ directions and effects among tissues suggests when pMEI affecting gene expression/splicing in multiple tissues, similar functional elements are affected. However, because gene expression and alternative splicing patterns are also correlated among related tissues (Fig. 1b, Fig. 3b), the similarity of eQTLs and sQTLs could also attribute to the correlated gene expression/splicing patterns among related tissues.

Although the QTL analyses can detect the association of pMEIs with gene expression and splicing changes, they do not provide information on the molecular mechanisms for the effect. By examining the enrichment of pMEIs, we found pMEIs in regions close to genes (intron, exon, 10 kb upstream or downstream) are more likely to correlate with gene expression and alternative splicing (Fig. 2, Fig S2). These pMEIs likely affect cis-elements (e.g., promoter, splicing sites, etc) of the associated genes. However, not all pMEIs identified in eQTL and sQTL analyses are near genes. Many of these pMEIs are far from the associated genes. These pMEIs may impact gene regulation through several mechanisms, such as serving as distal enhancers [44, 45], or altering chromatin looping structure [21, 37]. An interesting observation is that the effects of pMEIs on expression and splicing were highly correlated for some genes (Fig. 4). This may be because the regulation of gene expression was isoform-specific; the pMEI altered transcript level of specific isoforms and is then detected as both eQTL and sQTL. pMEIs with eQTL/sQTL signals are also highly enriched in enhancer regions (Fig. 2a, 2f). Because enhancers are key regulators for tissue-specific gene expression [46], this enrichment suggests that pMEIs could play a role in regulating tissue-specific expression and splicing.

In addition to the enrichment analysis, we also experimentally validate the predicted impact of several pMEIs using ectopic reporter assays. Such reporter assays are beneficial as several loci can be evaluated quickly to confirm computational predictions. However, while we have included as much of the endogenous locus as technically feasible, the in ectopic assay does not capture the full genomic context of the pMEI. Therefore, locus-dependent or tissue-specific effects may not be recapitulated in the reporter system. Further, the cloned pMEI locus were limited the subset of pMEIs we could evaluate. In the end, our experiments did validate the predicted effect of most of the tested pMEIs. To fully assess the functional impact of pMEIs, large scale functional validation will be needed in the future, including at the endogenous locus.

## Conclusions

Overall, our study showed that pMEIs are associated with thousands of gene expression and splicing variations in different tissues. Given the majority of the pMEI-associated eQTLs/sQTLs are tissue-specific and pMEIs are enriched in the enhancer regions, pMEIs could have a significant role in regulating tissue-specific gene expression/splicing. Detailed mechanisms for pMEIs’ role in gene regulation in different tissues will be an important direction for future studies.

## Method

### pMEI identification and filter

WGS data from the GTEx project v7 release were downloaded from dbGaP (phs000424.v7.p2). Of the 650 individuals in the v7 release, 12 were excluded from the analysis because of the issues during the dbGaP retrieval or the read mapping. WGS data from a reference sample HuRef (https://www.coriell.org/1/HuRef) was also included for quality control purposes. HuRef DNA sample was purchased from Coriell (NS12911, Camden, NJ, USA), and WGS was performed by Novogene (Sacramento, CA, USA) on the Illumina HiSeq platform using a PCR-free library and the Pair-End 150 bp sequencing format.

The Mobile Element locator Tool (MELT, version 2.1.5) [35] was used to identify pMEIs using the WGS data from the 639 individuals (638 GTEx individuals and HuRef). Briefly, WGS reads were aligned to the human reference genome GRCh38 with decoy sequence used in the 1KGP [47] using the Burrows-Wheeler Aligner (BWA, ver. 0.7.15) [48]. Output files were sorted and indexed with SAMtools (ver. 1.7) [49]. To identify pMEIs that are not present in the reference genome (nrMEIs), MELT (ver. 2.1.5) was run in the “MELT-SPLIT” mode under the default setting. The “MELT-SPLIT” mode includes five steps: Preprocess, IndivAnalysis, GroupAnalysis, Genotype, and MakeVCF. To identify pMEIs that are present in the reference genome but absent in the sequenced individuals (rMEIs), MELT was run in the “MELT-Deletion” mode which include two steps: Genotype and Merge. The ME reference files for *Alu*, LINE1, and SVA were downloaded within the MELT program. The final output is three files for nrMEIs and three for rMEIs in the VCF format.

The call sets were filtered to reduce false-positives and to focus on common variants. For nrMEIs, loci with <25% no-call rate, MELT ASSESS score ≥3, VCF filter column with “PASS” or “rSD”, and split reads >2 were kept. For rMEIs, sites with <25% no call rate were kept. For both nrMEIs and rMEIs, only loci with allele frequency between 0.05 and 0.95 in the dataset were kept. Hardy-Weinberg Equilibrium test were performed for each locus using individuals with “European” in the race description. Loci with p-value less than 10^−10^ is considered low-quality and were excluded from the analysis. The genomic coordinates of the loci were then lifted over from the human reference genome version GRCh38 to GRCh37/hg19 using CrossMap (ver. 0.2.7) [50]. Because of the known low-quality call on the Y chromosome, only loci from autosomes and X chromosome were kept for the downstream analysis.

### cis-eQTL mapping

Matrix eQTL (ver. 2.3) was used to identify association between genotypes and gene expression with a linear regression method [36]. Two genotype files were prepared: one file with only pMEIs for the ME-only analysis, and one file with pMEIs plus common SNPs and indels for the joint analysis. The SNP and indel genotypes were obtained from the GTEx project (phs000424.v7.p2, GTEx_Analysis_20160115_v7_WholeGenomeSeq_635Ind_PASS_AB02_GQ20_HETX_MISS 15_PLINKQC.PIR.vcf). The SNP and indels were filtered to remove sites with more than 25% no-call rate or with Hardy-Weinberg Equilibrium test p-value <10^−10^ in “European” individuals as described above.

Gene expression data in different tissues of different individuals were downloaded from GTEx website (https://gtexportal.org/home/datasets, GTEx_Analysis_2016-01-15_v7_RNASeQCv1.1.8_gene_tpm.gct.gz and GTEx_Analysis_2016-01-15_v7_RNASeQCv1.1.8_gene_reads.gct.gz). Normalized expression data of genes in each tissue were generated following the official GTEx QTL pipeline to reduce the effect of technical bias (https://github.com/broadinstitute/gtex-pipeline/tree/master/qtl). Briefly, in each tissue, a gene was kept if it has a TPM (Transcript Per Million) ≥ 0.1 and a raw read count ≥ 6 in ≥ 20% samples. Read counts among samples were normalized with the method described by [51] to obtain the trimmed mean of M values (TMM). Then, the TMM values of each gene were inverse normal transformed across the samples in each tissue.

The covariates for each tissue were downloaded from the GTEx website (https://gtexportal.org/home/datasets, GTEx_Analysis_v7_eQTL_covariates.tar.gz). The covariates include sex, three genotyping principal components, sequencing platform, and a various number of probabilistic estimation of expression residuals (PEER) factors based on the number of individuals (N) in each tissue type (15, 30, and 35 PEERs for N < 150, 150 ≤ N <250, N ≥ 250, respectively) [30, 52]. Input files for Matrix eQTL were generated with Python scripts for each tissues and Matrix eQTL were run with a window of 1 million bp (Mb) on either side of each gene. The p-value cutoffs (-p) were set at 1 for the ME-only analysis and 0.05 for the joint analysis. For the ME-only analysis, all genes were used as input and only eQTLs with FDR less than 10% by the Benjamini-Hochberg method were used for further analysis. For joint analysis, in each tissue, only genes reported in ME-only analysis with FDR < 10% were used as input for Matrix eQTL. From both eQTL analyses, a gene whose expression level showed an association with a variant with FDR < 10% in a given tissue is defined as an eGene. Protein-coding genes and non-coding genes are defined based on GENCODE gene models. Noncoding genes includes pseudogene, lincRNA, antisense, miRNA, misc_RNA, snRNA, snoRNA, rRNA, etc.

### cis-sQTL mapping

TPM values for each transcript and transcript models for each gene were downloaded from the GTEx website (https://gtexportal.org/home/datasets, GTEx_Analysis_2016-01-15_v7_RSEMv1.2.22_transcript_tpm.txt.gz and gencode.v19.transcripts.patched_contigs.gtf). ASEs were determined using SUPPA2 [53], with “–pool-genes” option enabled to group genes together if they are on the same genomic strand and share at least one exon. Seven types of ASEs were calculated: skipping exon (SE), alternative 5’ splice sites (A5), alternative 3’ splice sites (A3), mutually exclusive exons (MX), retained intron (RI), alternative first exons (AF), and alternative last exons (AL). Then, the PSI (percent spliced in) values were calculated by SUPPA2 based on the TPM values of transcripts in different tissues of different individuals. Similar to the eQTL analysis, sex, three genotyping principal components, sequencing platform, and PEER factors were included as covariates. PEER factors of different tissues were calculated by r-peer with PSI values [52]. The number of PEER factors were set based on number of individuals in each tissue type, same as in the eQTL analysis. ASEs with empty values were excluded as r-peer did not handle such cases. The cutoff for significant sQTLs was also set at 10% FDR.

### Fine mapping of causal variants for each eGene and ASE

CAVIAR (ver. 2.1) [54] was used to identify causal variants in the associated region for each eGene. CAVIAR takes a linkage disequilibrium (LD) file and a z-score file as inputs and reports a list of possible causal variants and the posterior probabilities of input variants being causal. pMEIs in the ME-only analysis and 100 most significant SNPs/indels in the joint analysis were chosen for each FDR-controlled eGene in the ME-only analysis. The signed r values for the LD file were calculated with PLINK (version 1.90) and the t-statistic values in Matrix eQTL output were used as the z-score. For each eGene, CAVIAR was run under the default setting (rho-prob 0.95, gamma 0.01, causal 1).

To identify causal cis-sQTL variants, similar analyses were performed as the eQTL analysis using CAVIAR (ver. 2.1). pMEIs in the ME-only analysis and 100 most significant SNPs/indels in the joint analysis were chosen for each FDR-controlled ASE in the ME-only analysis. Here, ASEs were used in place of eGenes, and PSI values were used in place of gene expression levels.

### Enrichment analysis of pMEIs

Fisher’s exact test was performed to check the enrichment of pMEIs in different regions of the affected genes. To test for enrichment in the eQTL analysis, common pMEIs were grouped into three categories based on their effect on gene expression: pMEIs not correlated with any gene (NS), correlated with at least one gene but not causal (Related), and being causal for at least one gene (Causal). For pMEIs grouped as NS and Related, the affected gene of a pMEI is defined as the gene with the smallest FDR value by Matrix eQTL no matter if the FDR is less than 10%; For causal pMEIs, the affected gene is the gene with pMEIs as causal variants and with the smallest FDR value. pMEIs that are not within 1 Mb window of any gene were excluded from the analysis. Functional genomic regions include enhancers from the Dragon Enhancers Database (DENdb, https://www.cbrc.kaust.edu.sa/dendb/src/enhancers.csv.zip) [55], 10 kb upstream from the transcription starting site (TSS), 10 kb downstream, exons, and introns of the affected gene. For each category, the number of pMEIs in different genomic functional groups were counted, and Fisher’s exact test was performed to determine the enrichment of pMEIs in those genomic regions in the Related and Causal categories relative to the NS category.

The enrichment analysis for sQTLs was performed similarly. For pMEIs grouped as NS and Related, the affected ASE of an ME is defined as the ASE with the smallest FDR value; for pMEIs grouped as Causal, the affected ASE is the ASE with pMEIs as causal variants and with the smallest FDR value. The affected gene is the gene contains the affected ASE. If ASE includes more than one gene, the longest gene was used to define the genomic functional groups. pMEIs that are not within 1 Mb window of any ASE were excluded from the analysis.

### Dual luciferase reporter assay for eQTLs

The effects of six representative pMEIs on gene expression were tested using a standard luciferase enhancer assay. For loci where the pMEI was predicted as causal for multiple eGenes the gene closest to the pMEI location was selected. About 300 bps of each genomic locus encompassing the pMEI insertion site were cloned into a modified pGL4.26 vector [56] using Gateway cloning (Invitrogen). The locus was amplified from 1KGP individuals using the primers in Table S4. For each locus, two independent clones were generated with the pMEI present and two clones without the pMEI. The orientation of the locus and the pMEI relative to the eGene was maintained relative to the luciferase reporter gene. All constructs were verified by Sanger sequencing. The firefly luciferase vectors were each co-transfected with a Renilla plasmid (pRL, Promega) into 293T cells using Fugene HD (Promega). After 48 hours, luciferase levels were measured using Dual-glo luciferase assay system (Promega) and the GloMax-Multi Detection System (Promega). Firefly and Renilla levels were normalized to background in wells with no transfected plasmids and a ratio of firefly to renilla levels in each well accounted for any differences in transfection efficiency. Results were graphed as relative luciferase units for each construct and an un-paired 2-tailed t-test was performed for each locus.

### Ectopic minigene reporter assay for sQTLs

The effects of four representative pMEIs on alternative splicing were experimentally evaluated with an ectopic minigene reporter assay as previously described [14]. Briefly, for each locus, a genomic fragment surrounding the pMEI and nearby exons was cloned into an intron between rat insulin exons in the pSpliceExpress vector (Addgene) [57] using Gateway cloning (Invitrogen). The region was amplified using primers listed in Table S4 from DNA of 1KGP individuals. Two constructs were generated for each evaluated locus: one with the pMEI present and one without the pMEI. Two independent clones were isolated for each construct and verified by Sanger sequencing. The plasmids were transfected (Fugene HD, Promega) into 293T cells and after 24 hours RNA was extracted (Quick RNA MicroPrep Kit, Zymo Research) and reverse transcribed to cDNA (iScript cDNA Synthesis Kit, BioRad). RT-PCR was performed with primers that bind within the rat insulin exons (Ins1: 5′-CAGCACCTTTGTGGTTCTCA-3′ and Ins2: 5′-AGAGCAGATGCTGGTGCAG-3′). For the *IFT122* locus, to enable a sensitive quantification of the rare alternative exon, we increased specificity by repeating the RT-PCR with a primer in the constitutive exon from this locus (5’-AAAGTAAAGATCGAGCGGCC-3’ paired with Ins2). For each locus, the relative quantification of alternatively spliced RNA isoforms was performed on ethidium bromide stained agarose gels with band intensities normalized for DNA fragment length. Two transfections were performed for each independent clone of each construct, resulting in four data points for each type of construct (i.e., with or without the pMEI) for each locus. Quantification is graphed as percent of transcripts that include the alternative exon and unpaired t-tests compared the percent inclusion when the pMEI was present versus absent at each locus.

## Abbreviations

MEs: Mobile elements
pMEIs: Polymorphic mobile element insertions
GTEx: Genotype-Tissue Expression
QTL: Quantitative trait loci
eQTLs: Expression quantitative trait loci
sQTLs: Splicing quantitative trait loci
SNPs: Single nucleotide polymorphisms
LTR: Long terminal repeat
SINE: Short interspersed element
LINE1/L1: Long interspersed element 1
SVA: SINE-VNTR (variable-number tandem repeat)-Alu
1KGP: The 1000 Genomes Project
WGS: Whole genome sequencing
MELT: Mobile Element locator Tool
BWA: Burrows-Wheeler Aligner
nrMEIs: Non-reference mobile element insertions
rMEIs: Reference mobile element insertions
VCF: Variant call format
PEER: Probabilistic estimation of expression residuals
TMM: Trimmed mean of M values
FDR: False discovery rate
ASEs: Alternative splicing events
SE: Skipping exon
A5: Alternative 5’ splice sites
A3: Alternative 3’ splice sites
MX: Mutually exclusive exons
RI: Retained intron
AF: Alternative first exons
AL: Alternative last exons
PSI: Percent spliced in
LD: Linkage disequilibrium
DENdb: Dragon Enhancers Database
SD: Standard deviation
TPM: Transcript per million
LCLs: Lymphocyte cell lines

## Declarations

### Ethics approval and consent to participate

Not applicable.

### Consent for publication

Not applicable.

### Availability of data and materials

The VCF files of individual pMEI genotypes are available under dbGaP project “Impact of Mobile Element Insertions on Human Transcriptome Variation” (Study ID: 38256).

### Competing interests

The authors declare no competing interests.

### Funding

This study was supported by the startup fund from the Human Genetics Institute of New Jersey to JX.

### Authors’ contributions

XC and JX designed the research. The analysis was performed primarily by XC and YZ, and some by HL and JX. The validation experiments were performed by LP, JS, and KB. XC, YZ, LP, and JX wrote the draft of the manuscript. All authors read and approved the final version of the manuscript.

## Acknowledgement

The Genotype-Tissue Expression (GTEx) Project was supported by the Common Fund of the Office of the Director of the National Institutes of Health, and by NCI, NHGRI, NHLBI, NIDA, NIMH, and NINDS. The data used for the analyses described in this manuscript were obtained from dbGaP accession number phv00169064.v7.p2 on February, 2018.

## Additional files

**Additional file 1: Supplemental Figures**

**Fig. S1.**
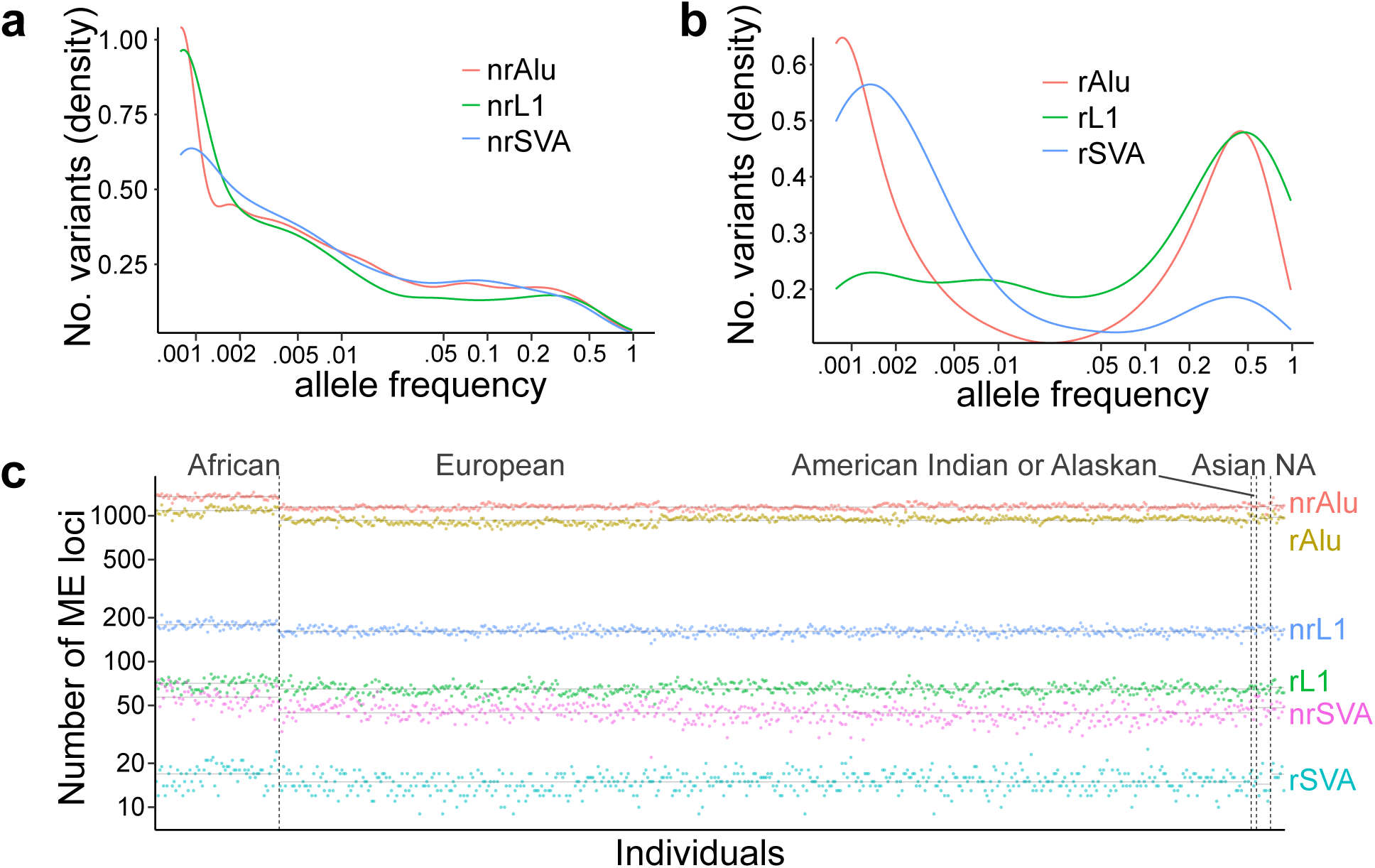
Overview of pMEIs in GTEx individuals. Raw output of MELT was filtered as described in methods to include only high confidence loci. (**a, b**) Allele frequency distribution of nrMEIs (**a**) and rMEIs (**b**). (**c**) Counts of nrMEIs (nrAlu, nrL1, nrSVA) and rMEIs (rAlu, rL1, rSVA) relative to the reference genome (GRCh38) in each individual. Individuals are grouped based on ethnic groups. NA, ethnic group unavailable.

**Fig. S2.**
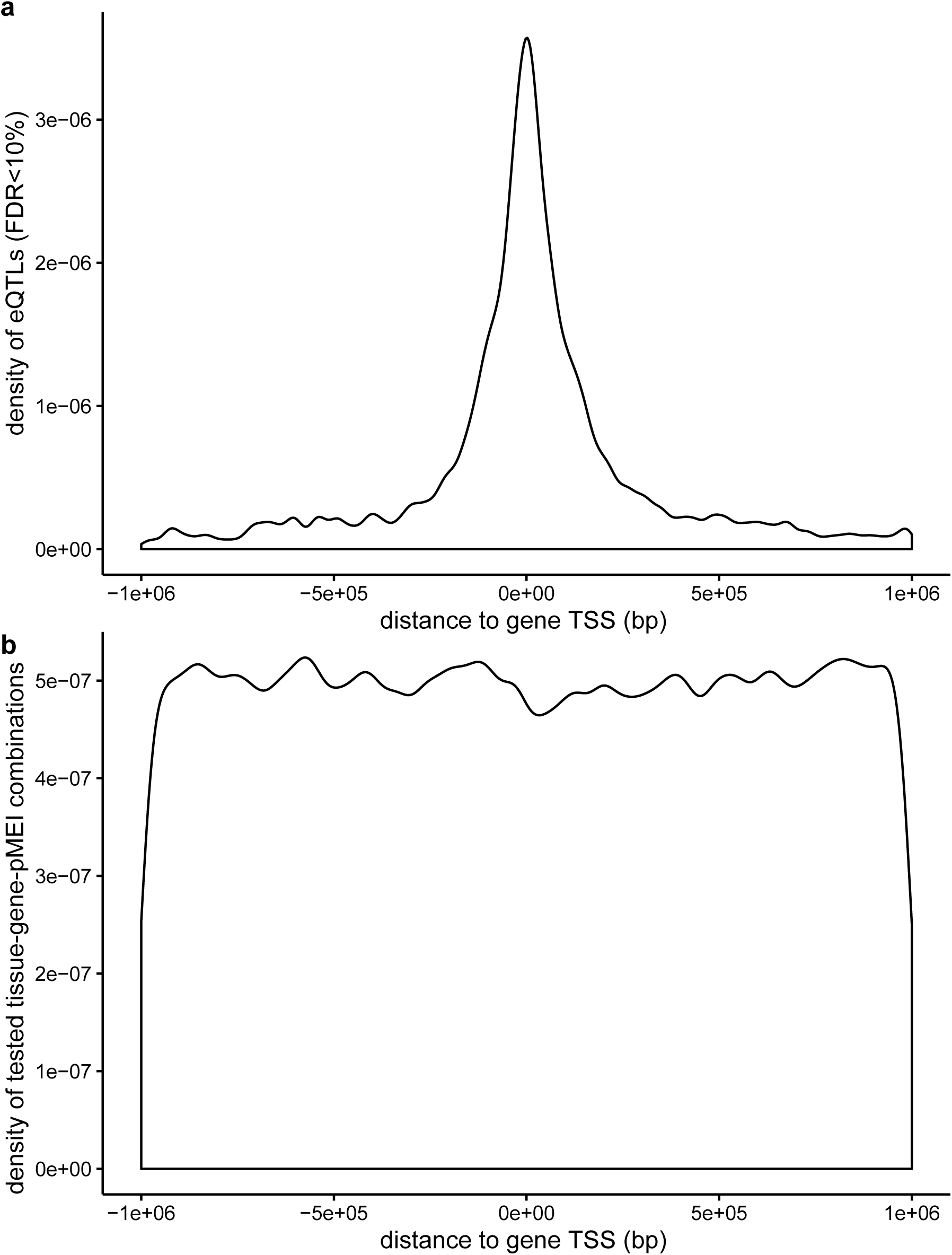
Enrichment of pMEIs around Transcription Starting Sites (TSSs) of genes. **(a)** Density of pMEIs-associated with eQTLs (tissue-gene-pMEI combinations with FDR < 0.1) around TSSs. **(b)** Density of pMEIs in all possible tissue-gene-pMEI combinations examined by Matrix eQTL around TSSs.

**Fig. S3.**
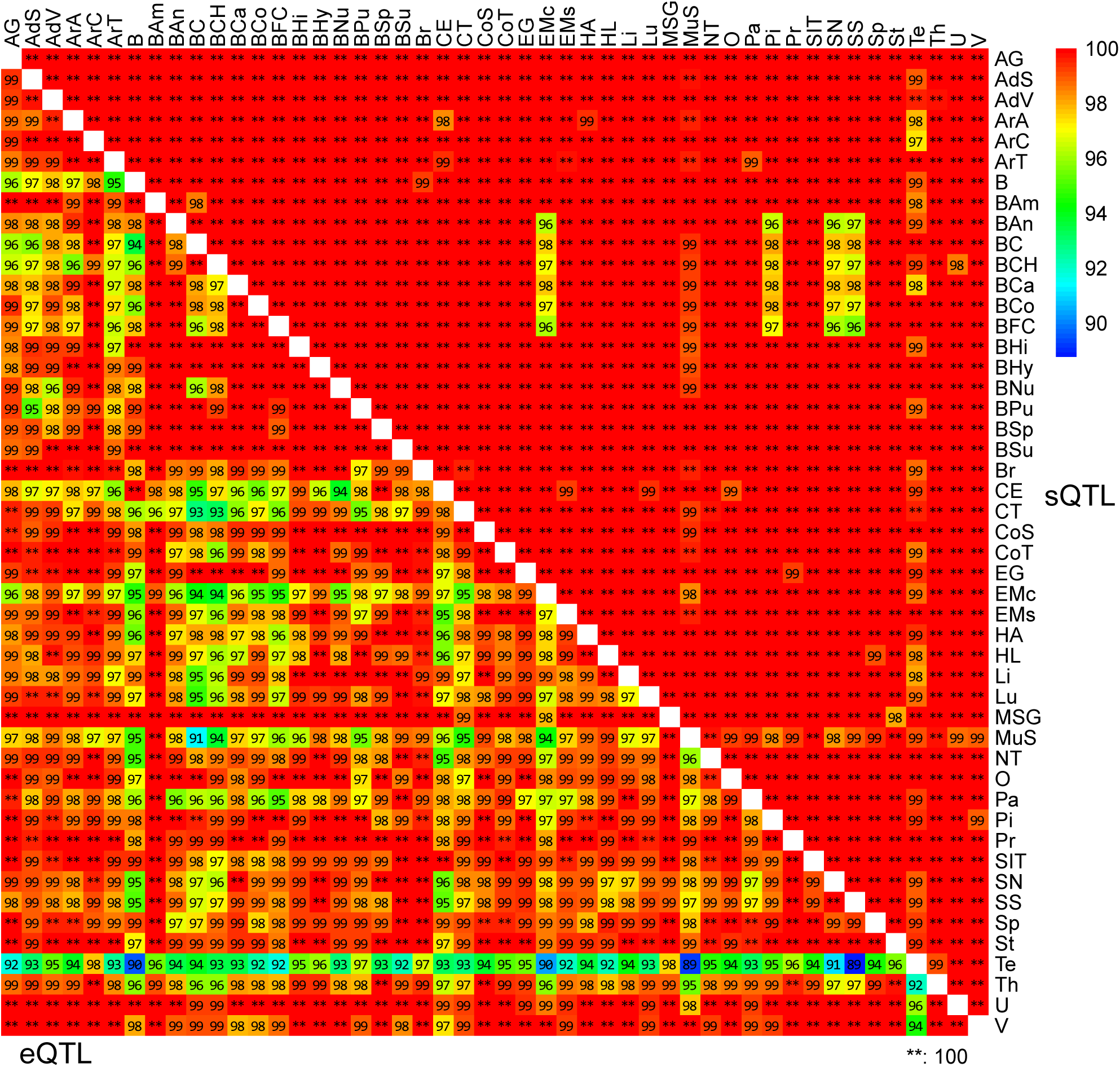
Agreement of the effect direction between a pair of tissues for eQTLs (lower-left) and sQTLs (upper-right). Numbers are the percentage of shared eQTL/sQTL of two tissues with the same impact direction (the sign of beta value). Tissue abbreviations were the same as in Fig. 1. **=100% shared.

**Additional file 2: Supplemental Tables**

**Table S1: Number of samples, expressed genes, and eQTL results for each tissue**.

**Table S2: Number of samples and sQTL results for each tissue**.

**Table S3: LCL eQTLs in the current study and an 1KGP study (Spirito et al. 2019)**.

**Table S4: Primers for eQTL and sQTL cloning**.

## Notes

### Competing Interest Statement

The authors have declared no competing interest.

